# Heterochronic Developmental Shifts Underlie Floral Diversity within *Jaltomata* (Solanaceae)

**DOI:** 10.1101/169060

**Authors:** Jamie L. Kostyun, Jill C. Preston, Leonie C. Moyle

## Abstract

**Background:** Heterochronic shifts during mid to late stages of organismal development have been proposed as key mechanisms generating phenotypic diversity. To determine whether late heterochronic shifts underlie derived floral morphologies within *Jaltomata* (Solanaceae) – a genus whose species have extensive and recently evolved floral diversity – we compared floral development of four diverse species (including an ambiguously ancestral or secondarily derived rotate, two putatively independently evolved campanulate, and a tubular morph) to the ancestral rotate floral form, as well as to an outgroup that shares this ancestral floral morphology.

**Results:** We determined that early floral development (<1 mm bud diameter, corresponding to completion of organ whorl initiation) is very similar among all species, but that different mature floral forms are distinguishable by mid-development (>1 mm diameters) due to differential growth acceleration of corolla traits. Floral ontogeny among similar mature rotate forms remains comparable until late stages, while somewhat different patterns of organ growth are found between species with similar campanulate forms.

**Conclusions:** Our data suggest shared floral patterning during early-stage development, but that different heterochronic shifts during mid- and late-stage development contributes to divergent floral traits. Heterochrony thus appears to have been important in the rapid and repeated diversification of *Jaltomata* flowers.

## Background

In The Origin of Species, Darwin stated “we can actually see in embryonic crustaceans and in many other animals, and in flowers, that organs, which when mature become extremely different, are at an early stage of growth exactly alike.” [1]. This prescient statement both suggests the importance of understanding diversity in a developmental and phylogenetic framework [2], and raises the question about which mechanisms underlie these hypothesized mid to late stage shifts in ontogeny. Heterochrony—a change in the relative timing or rate of a developmental process between a derived lineage and its ancestor—is one potential mechanism contributing to such trait variation [3-4]. Since being formally defined by Haeckel [5], the concept of heterochrony has undergone significant changes in its usage and application [6-9; reviewed in 10-11]. Despite these differences in the exact definition utilized, heterochrony is still broadly viewed as an important framework in which to examine and understand morphological changes [3; 10], especially in animal taxa. For instance, the classic example of axolotls that retain more juvenilized features than their ancestors [12], the large variation in cranial morphology among different dog breeds [13], and even the evolution of extremely complex traits such as metamorphosis in insects [14], have largely been attributed to heterochronic shifts during development.

The general concept of heterochrony, as well as the distinct processes that comprise this mechanism of developmental change, can also be meaningfully applied to plant evolution [15-16]. Heterochrony is most often identified in plants by examining individual organs or functional units, such as flowers or individual leaves [4]. Broadly, heterochronic shifts include changes in the relative timing of initiation (i.e. onset) or termination (i.e. offset) of a developmental process – which together determine the duration of this developmental process. Additionally, changes can occur in the rate at which a developmental process proceeds. Shorter developmental duration or decreased growth rate (termed pedomorphosis) often results in the reduction or juvenilization of a trait, while longer developmental duration or increased growth rate (termed peramorphosis) often results in the elaboration of a trait. Both types of shifts – duration or rate, as well as their combined effects – can conceivably contribute to observed variation in plant traits [3-4, 17].

Unlike plant architecture in general, flowers undergo determinate development, and thus are particularly attractive subjects for studying the contribution of heterochrony to morphological divergence. While most flowers conform to a shared ground plan, i.e. egg and pollen-producing structures surrounded by a non-reproductive perianth (sepals and petals), many lineages exhibit abundant inter- and intra-specific variation in the organization (phyllotaxy), number (merosity), size and shape, degree of fusion, and specific identity of floral organs [18-19]. Numerous studies have examined the potential role of heterochronic changes in floral trait evolution, including those associated with mating system transitions. The shift from outcrossing to predominately selfing is considered one of the most common evolutionary transitions among plants [20-21], and is typically associated with reduction of flower size, petal size, and/or anther-stigma separation (herkogamy) – collectively referred to as the ‘selfing syndrome’ [22-23]. Comparative work examining closely related outcrossing and selfing species, as well as among intra-specific populations [24], has generally revealed that reductions in floral size result from pedomorphic changes, either from decreased growth rates (i.e. neoteny; e.g. [25]), or truncation of the growth period [26]. However, some studies have also found that smaller flowers or those with reduced herkogamy in selfing lineages result from an increased growth rate over either short [27] or long [28] developmental periods, as well as different growth patterns between populations with high selfing rates [24]. Similarly, comparative work examining the developmental basis of cleistogamous flowers (where sexual maturation and self-fertilization occurs within closed buds) also indicates a prominent role of heterochronic changes [29]. Such cleistogamous flowers have been found to result from various combinations of heterochronic shifts, including decreased organ growth rates and shortening of the duration of development (e.g. [30]), or decreased growth rates of petals and stamens (e.g. [31]). In contrast however, Luo et al. [32] determined that cleistogamous flowers in *Pseudostellaria heterophylla* result from induction of fewer organ primordia (including complete loss of petal primordia initiation) than in chasmogamous (i.e. open flowers) on the same individual. Thus, several distinct heterochronic (and in fewer reported cases, non-heterochronic) processes can result in convergent floral phenotypes associated with mating system transitions, such as selfing syndromes and cleistogamy.

Apart from developmental studies of mating system transitions, comparative ontogenetic studies have examined floral trait variation likely related to pollinator shifts [33-35]. A classic example is Guerrant’s [33] examination of developmental differences between a pair of *Delphinium* species – one bee-pollinated, the other hummingbird-pollinated. In that instance, the derived hummingbird-pollinated flower results from an overall decreased growth rate (pedomorphosis via neoteny), but also accelerated growth for an extended period (peramorphosis via acceleration and hypermorphosis) specifically in the reward-providing nectariferous petals. More recently, Armbruster et al. [35] described the evolution of buzz-pollinated flowers in *Dalechampia* via neoteny, with sexual maturity of the anthers occurring at an earlier developmental stage (i.e. similar to pre-anthesis ancestral buds) than in species with the post-anthesis ancestral floral form (i.e. with opened and extended petals and sepals). Nonetheless, despite this increasing attention to the role of heterochrony in plant development, several critical questions remain, including how frequently heterochronic shifts underlie changes in floral form, and whether convergent phenotypes may be the result of similar or divergent developmental mechanisms.

*Jaltomata* (Solanaceae) includes approximately 60-80 species distributed from the Southwestern United States to the Andean region of South America, with extensive and recently evolved floral diversity [36-39]. Although the pollination biology of this genus has yet to be formally evaluated, field observations (T. Mione and S. Leiva G., pers. comm.) reveal that different species are visited by bees and hummingbirds. In addition, *Jaltomata* produce varying amounts of floral nectar as a pollinator reward, unlike close relatives like *Solanum* [40-41]. Examination of corolla morphology variation within Solanaceae [41] reveals that the closest lineages to *Jaltomata* (*Solanum*, *Capsicum*, and *Lycianthes*) all have predominately flattened ‘rotate’ corollas, suggesting that this rotate form is ancestral within *Jaltomata*. Indeed, ancestral state reconstruction on a molecular phylogeny of 45 *Jaltomata* species [37] supports the rotate corolla form as ancestral, and that both bell-shaped ‘campanulate’ and elongated ‘tubular’ flowers are derived specifically within the genus (Fig. 1). It also suggests that several instances of the independent evolution of either derived campanulate or tubular floral forms in different *Jaltomata* species; in contrast, both campanulate and tubular forms generally appear to produce more nectar than rotate forms ([42]; T. Mione, pers. comm.; J. L. Kostyun, unpublished). Phylogenetic reconstruction with fewer species but additional loci [36, 38], as well as phylogenomic reconstruction using whole transcriptomes (Wu, Kostyun, Moyle, unpublished), also recover the major clades identified in [37] further supporting the inferences that rotate corolla morphology is ancestral within the genus and that campanulate corollas in different clades are likely convergent rather than homologous.

**Fig. 1.**
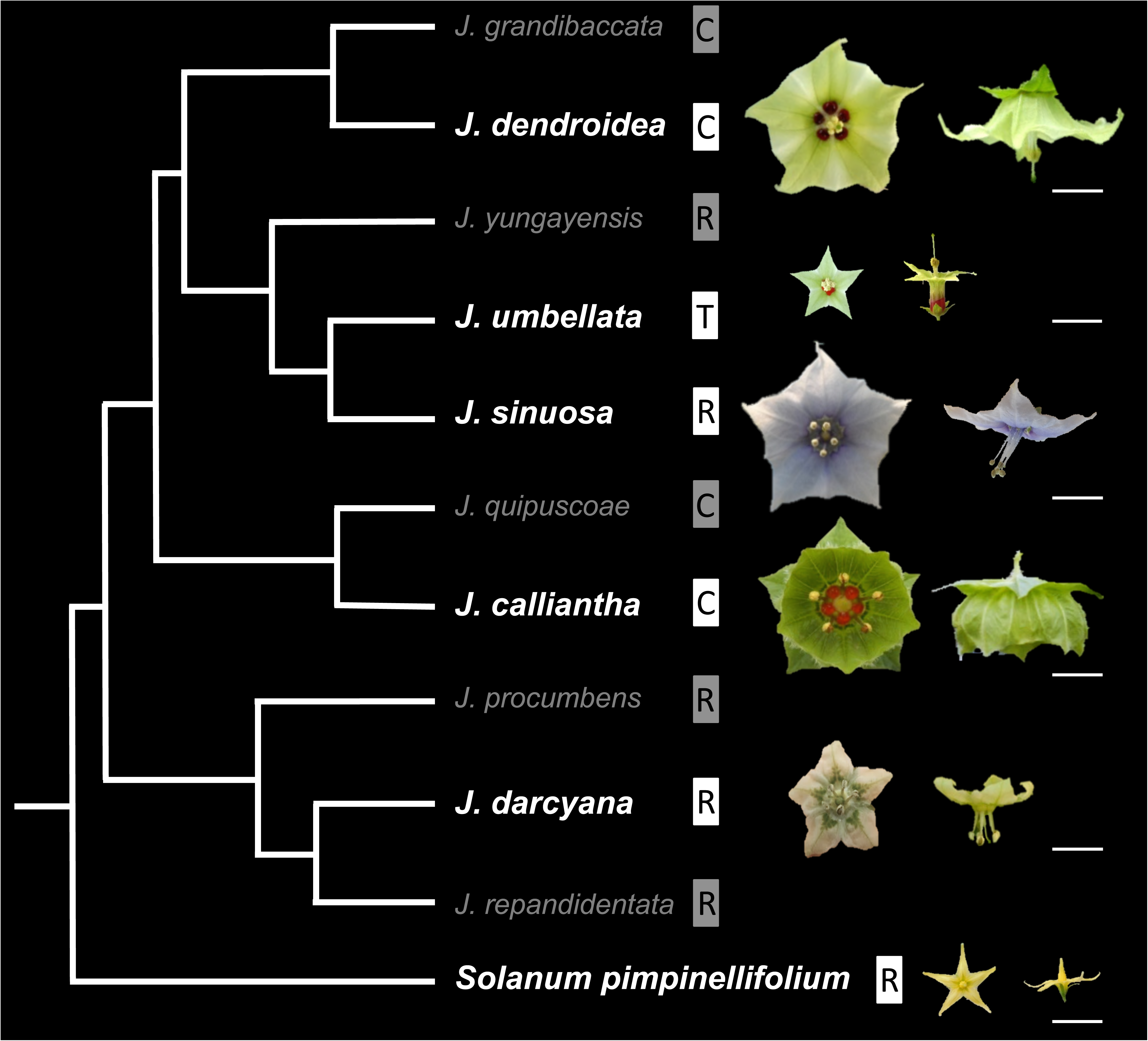
Simplified phylogenetic relationships in *Jaltomata* and the outgroup *Solanum pimpinellifolium* based on [38, 41] and (Wu, Kostyun, Moyle, unpublished). Representative mature flowers are depicted for the species examined here (in bold font). White scale bars = 1 cm. R, rotate; C, campanulate; T, tubular. Note: All five focal *Jaltomata* species are represented in [37], but as this study was accepted prior to formal naming of *J. calliantha* [56], this species is called “J. hummingbird”.

We hypothesized that both derived campanulate and tubular forms might represent elaborated versions of the ancestral rotate form, specifically through variation in corolla (petal) growth prior to flower anthesis (opening). We therefore assessed whether heterochronic changes (specifically peramorphic changes such as longer duration of development and/or growth acceleration, which often produce larger or more elaborated structures) underlie these transitions, particularly during late stages of ontogeny [1-2, 4]. (Therefore, although heterochrony can also refer to changes in the actual developmental events (i.e. sequence heterochrony; [11]), here we are particularly interested in assessing changes in growth). In addition to evaluating the role of heterochrony in the evolution of derived floral forms, we also expected that comparing floral development between species with similar mature morphs--but unresolved phylogenetic relatedness--would provide insight into their evolutionary origin [37] (Wu, Kostyun, Moyle, unpublished). For instance, if similar mature forms result from similar developmental processes, this could suggest re-use of similar pathways or a shared single origin. In contrast, if they result from distinct developmental processes, this could indicate that these forms are independently derived, or that these forms share a common evolutionary origin (i.e. are homologous) but underwent developmental systems drift [43] during their divergence.

Given these considerations, our main goals in this study were to: 1) assess evidence for heterochronic changes during floral development in species with derived campanulate and tubular forms; 2) determine which type(s) of heterochronic changes are associated with specific floral trait changes; and 3) assess whether putatively convergent forms might result from similar or different developmental processes (i.e. are associated with the same or different types of heterochronic shifts). To address these goals, we compared floral ontogeny of four *Jaltomata* species with divergent corolla morphologies to a *Jaltomata* species with a rotate corolla that is representative of the inferred ancestral state (i.e. plesiomorphic), as well as an outgroup species that also has an inferred plesiomorphic rotate corolla form.

Our findings support the inference that a combination of parallel and convergent allometric changes (i.e. shifts in the size of particular floral organs in relation to the entire flower) have given rise to floral variation in this group, and show that heterochronic shifts explain these changes. In particular, two types of peramorphic changes (extended duration of floral development and accelerated growth) predominately explain changes in corolla traits. Thus, peramorphism emerges as an important developmental mechanism controlling diversity of *Jaltomata* corolla forms. Finally, species with similar mature campanulate corollas follow similar but clearly not identical growth trajectories, consistent with phylogenetic inferences that these campanulate floral morphs have independent origins [37]; (Wu, Kostyun, Moyle, unpublished) (Fig. 1). In contrast, we found that species with similar mature rotate corollas follow nearly identical developmental trajectories (specifically, that overall bud and organ growth rates do not differ) until the very last stages of ontogeny, suggesting a single evolutionary origin of the rotate form in this case.

## Methods

### Plant materials and growth conditions

Field-collected seeds of our five target *Jaltomata* species were provided by Dr. Thomas Mione (Central Connecticut State University), and seed for one wild tomato outgroup (*Solanum pimpinellifolium*, accession LA1589) was obtained from the Tomato Genetics Resource Center at the University of California, Davis (Table S1). We selected this outgroup because *Solanum* is considered the sister genus to *Jaltomata* [36, 38], this species has an ancestrally representative rotate corolla [41], floral development in this species has previously been characterized [26, 44], and--as a wild species--its floral development has not been influenced by domestication (i.e. compared to domestic tomato, *S. lycopersicum*). Our focal *Jaltomata* species included the Central American species *Jaltomata darcyana* that has an rotate corolla representative of the inferred ancestral corolla morphology, and four species found as natives exclusively in South America: *Jaltomata calliantha* and *Jaltomata dendroidea* with putatively independently derived campanulate corollas, *Jaltomata umbellata* with a tubular corolla, and *Jaltomata sinuosa* with a rotate corolla that is ambiguously ancestral or secondarily derived (Fig. 1). Plants were cultivated in the Indiana University Research Greenhouse, under standardized temperature (15-20°C) and light (16-hour days) conditions. All plants were watered to field capacity on automatic driplines, and received weekly fertilizer treatment.

### Floral development

To estimate the duration of floral development and overall absolute floral growth rate, young inflorescences were first identified when the largest bud was approximately 1 mm in diameter, and then evaluated every second day until first anthesis (petal opening). In particular, we evaluated and measured bud diameter on the largest (oldest) bud from at least three inflorescences on three individuals per species, except for *J. dendroidea* for which we could only measure buds from two individuals (three and two inflorescences per individual, respectively).

To assess relative organ growth during development, buds up to 4 mm in diameter were collected into 70% ethanol or formalin-acetic acid-alcohol (FAA), dissected as required, dehydrated through an ethanol series at room temperature, and incubated in a 4°C fridge overnight. Samples were critical point dried (using a Balzers CPD030), mounted on aluminum stubs, and sputter-coated with gold-palladium (using a Polaron E5100). They were then imaged using a JEOL 6060 or a JEOL 5800LV scanning electron microscope (SEM) at the University of Vermont or Indiana University, respectively. Morphological traits were measured on the resulting bud micrographs using ImageJ [45]. For buds greater than 4 mm in diameter, morphological traits were measured by hand on freshly collected samples using digital calipers. To ensure that we were comparing organ sizes at the same developmental stage across species, we assigned measured buds to discrete developmental stages based on morphological markers, following [44] for which floral developmental stages were determined in the same accession of *S. pimpinellifolium* as used here. This resulted in six distinct ‘early’ developmental stages, one ‘mid’ stage, and three ‘late’ stages, based on these morphological markers (Table 1).

**Table 1.**
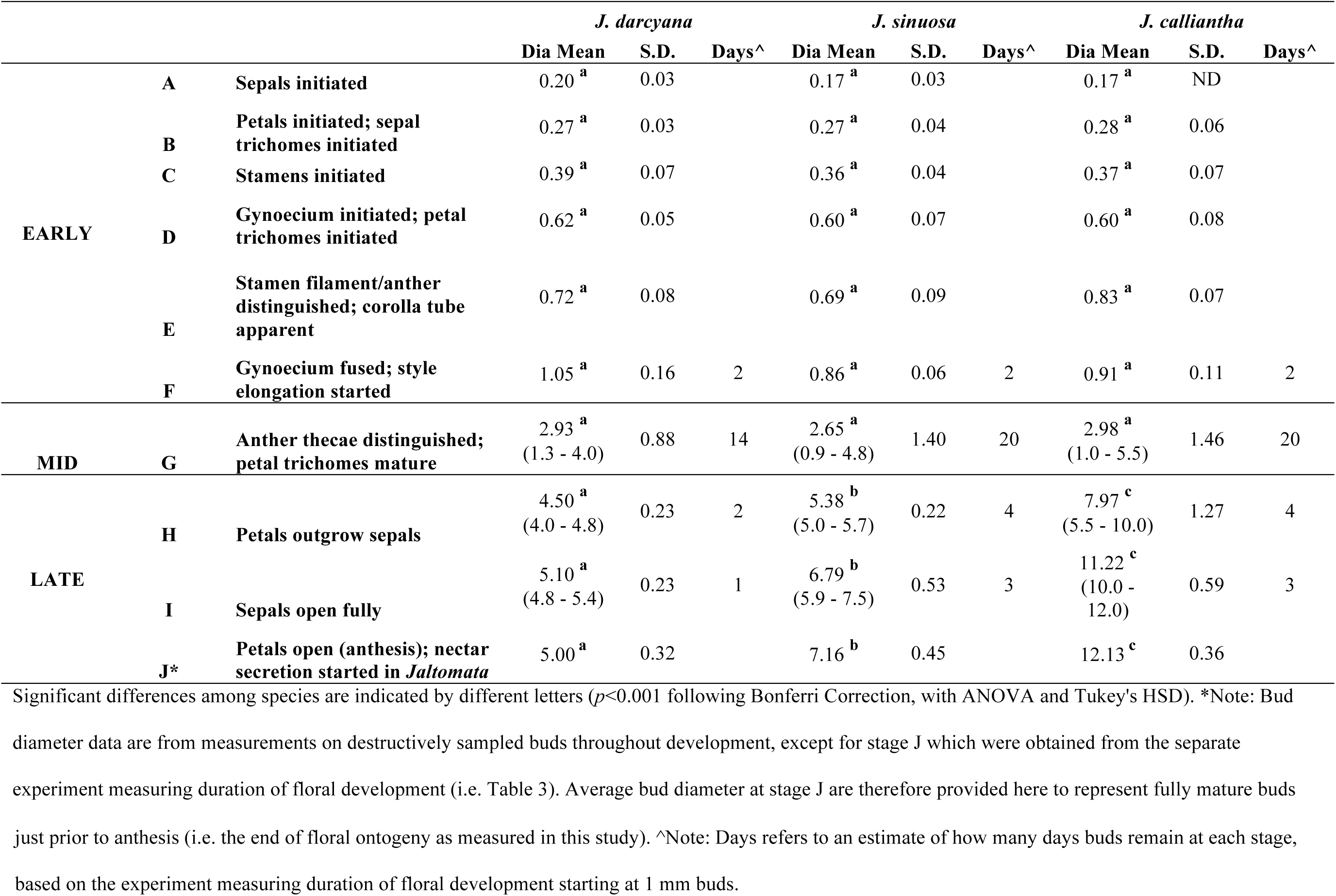
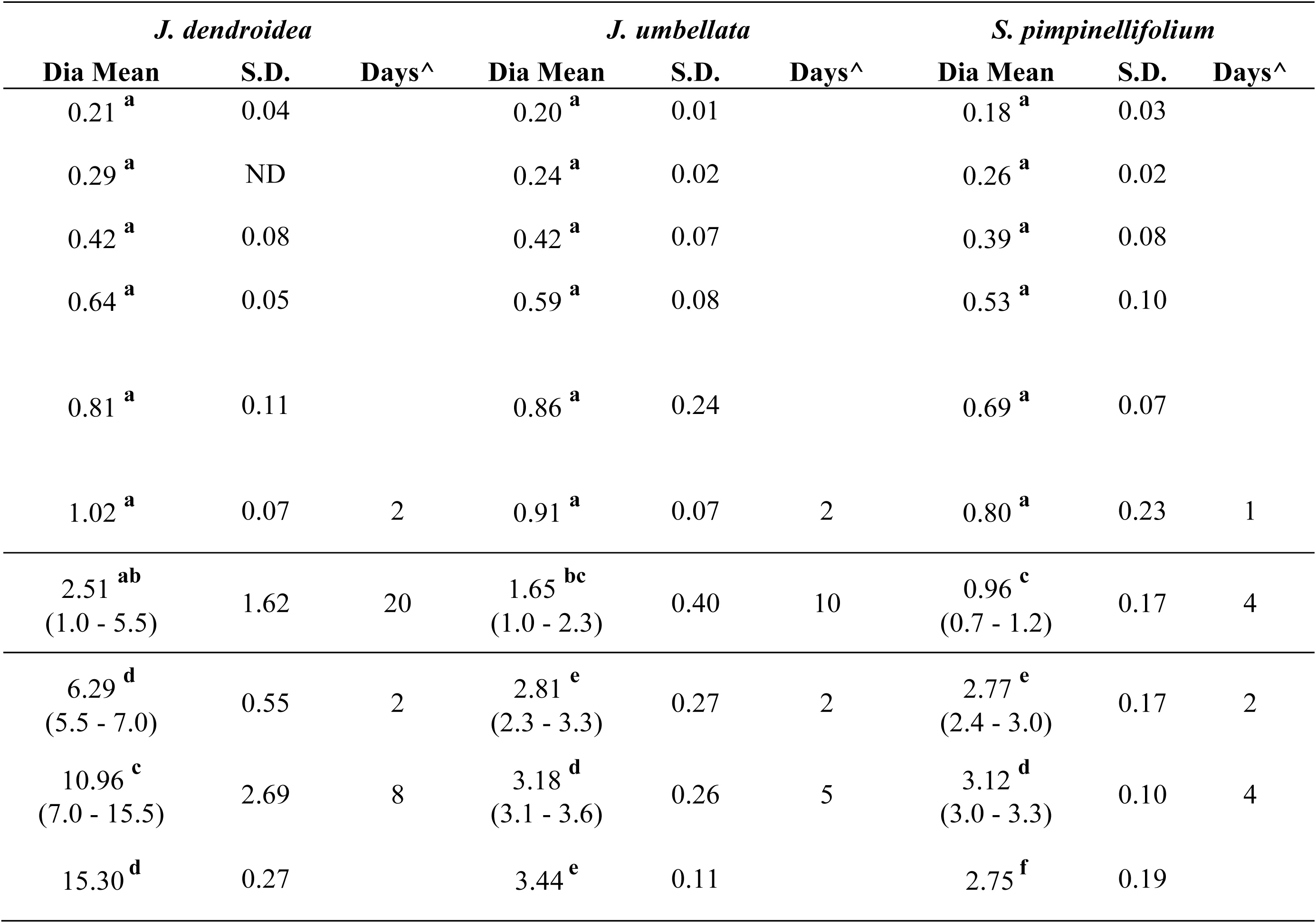
Bud development stages based on morphological markers, means (and ranges, in parentheses, for mid- and late stages) of bud diameter for each stage across species, and estimates of duration (in days) spent in each stage.

Given the potential role of trichomes in regulating corolla shape during bud development [46], we also determined the developmental timing of trichome initiation and maturation. In particular, we assessed whether divergent corolla morphologies are associated with differences in trichome patterning during floral development. Because our focal species also vary in nectar volume per flower, and derived campanulate and tubular forms are generally associated with greater nectar volume across the genus (T. Mione, pers. comm.; J. L. Kostyun, unpublished), we determined the onset of nectar secretion using a 20x hand-lens. Our primary goal was to determine whether this observed difference in nectar volume between divergent corolla forms is associated with changes in the developmental timing of nectar secretion.

### Mature floral trait measurements

Mature floral traits were measured with hand-held digital calipers on three flowers each for at least three individuals per species, and included calyx diameter, sepal length, corolla diameter, corolla depth, corolla fusion (i.e. length of the corolla tube), petal length, lobe length, stamen length, anther length, ovary diameter, and style length (Fig. S1). To account for potential size differences among flowers within an inflorescence, floral traits were only measured on the oldest flower within each examined inflorescence. For *Jaltomata* species only (the outgroup does not produce nectar), we also measured nectar volume per flower to the nearest 1 µL with a pipette. To reduce potential environmental effects on nectar production, nectar volume per flower was always measured during the early afternoon following watering. Trichome type(s) present on sepals and petals was also scored using a 20x hand lens.

### Statistical analyses

We used analyses of variance (ANOVA), followed by Tukey’s honest significant difference (HSD) post-hoc tests, to assess differences among species in three comparisons: on mature floral traits; log-transformed floral organ size at discrete stages during development (as specified in Table 1); and duration (i.e. number of days) of floral development.

To compare relative floral organ growth rates during development, we used linear regression on log-transformed bud measurements. Because buds grow at different rates among species (Table 1), we regressed organ sizes on bud diameter to account for overall floral size; thus, we assessed whether there were significant interaction effects between bud diameter and species as a measure of differential growth rates. We performed these analyses in two ways. First, we used data from the entire course of floral development. However, because we determined that species differences in bud growth (specifically, bud diameter) only occur once all floral organs are initiated (i.e. starting in stage G, Table 1), we also performed these regressions separately for two discrete groups of developmental stage (“Early” versus “Mid” + “Late”, in Table 1). This enabled us to more directly examine organ growth in relation to overall bud growth; we present the latter set of analyses in the main text, and the former (i.e. over total floral development) within Table S3 for comparison. Because we are most interested in potential heterochronic shifts within *Jaltomata*, we used *J. darcyana* as the baseline in these analyses to assess differences between ancestral and derived floral forms. However, since *J. darcyana* flowers could have somewhat derived developmental trajectories (and the actual ancestor of all five *Jaltomata* species is no longer extant), we also performed these analyses using either *S. pimpinellifolium* (as another representative of the inferred rotate ancestral state) or *S. pimpinellifolium* + *J. darcyana* (as a ‘composite’ representation of the inferred rotate ancestral state – specifically, we treated these species as a single group) as the baseline. These analyses returned qualitatively consistent results as our comparisons using *J. darcyana* (see Results), and are provided in Table S3 for comparison. We also compared organ growth rates between the two derived campanulate forms (i.e. *J. calliantha* versus *J. dendroidea*). All analyses were performed within the R statistical environment [47].

## Results

### Mature floral traits differ markedly across species

Mature, post-anthesis *Jaltomata* flowers differ substantially in multiple traits, including overall size and relative sizes of individual organs, volume of nectar produced per flower, and trichome type and density on sepals and petals (Table 2; Fig. 1). In terms of floral size, campanulate *Jaltomata calliantha* and *J. dendroidea* have the largest flowers, while rotate *S. pimpinellifolium* has the smallest; *Jaltomata* flowers also differ from *S. pimpinellifolium* in having free stamens (compared to a fused anther cone, considered a derived trait in the wild tomatoes [41]), and in producing nectar. Calyx diameter significantly differs across all species, while both corolla and ovary diameter significantly differ between all species pairs, except tubular *J. umbellata* versus *S. pimpinellifolium* (ANOVA *p* < 0.00001; Tukey HSD, *p* < 0.0001), which both have comparatively small flowers. Although mature floral organ sizes vary substantially among *Jaltomata* species, the ancestrally rotate form of *J. darcyana* flowers have the shortest petals (ANOVA *p* < 0.00001; Tukey HSD, *p* < 0.0001), and the smallest amount of corolla fusion (ANOVA *p* < 0.00001; Tukey HSD, *p* < 0.00001) (Table 2).

**Table 2.** Mature floral trait means for the five included *Jaltomata* species and *Solanum pimpinellifolium*.

|  | *J. darcyana*<br>(n=7) |  | *J. sinuosa*<br>(n=7) |  | *J. calliantha*<br>(n=7) |  | *J. dendroidea*<br>(n=3) |  | *J. umbellata*<br>(n=5) |  | *S. pimpinellifolium*<br>(n=3) |  |
| --- | --- | --- | --- | --- | --- | --- | --- | --- | --- | --- | --- | --- |
|  | Mean | S.D. | Mean | S.D. | Mean | S.D. | Mean | S.D. | Mean | S.D. | Mean | S.D. |
| **Calyx Diameter (mm)** | 11.88 <sup>a</sup> | 0.37 | 15.49 <sup>b</sup> | 0.40 | 33.88 <sup>c</sup> | 1.21 | 19.69 <sup>d</sup> | 1.74 | 8.14 <sup>e</sup> | 1.63 | 5.77 <sup>f</sup> | 0.43 |
| **Sepal Length (mm)** | 6.32 <sup>a</sup> | 0.16 | 7.48 <sup>b</sup> | 0.25 | 18.01 <sup>c</sup> | 1.10 | 9.67 <sup>d</sup> | 0.80 | 4.41 <sup>e</sup> | 0.73 | 3.45 <sup>f</sup> | 0.23 |
| **Corolla Diameter (mm)** | 20.48 <sup>a</sup> | 2.18 | 29.82 <sup>b</sup> | 2.27 | 27.48 <sup>c</sup> | 3.48 | 35.35 <sup>d</sup> | 3.27 | 15.32 <sup>e</sup> | 2.52 | 15.58 <sup>e</sup> | 0.58 |
| **Corolla Depth (mm)** | 0.77 <sup>a</sup> | 0.09 | 1.38 <sup>a</sup> | 0.27 | 15.57 <sup>b</sup> | 1.84 | 12.72 <sup>c</sup> | 0.88 | 10.25 <sup>d</sup> | 2.27 | 0.45 <sup>a</sup> | 0.07 |
| **Corolla Fusion (mm)** | 3.96 <sup>a</sup> | 0.28 | 9.45 <sup>b</sup> | 0.46 | 15.64 <sup>c</sup> | 0.44 | 14.91 <sup>c</sup> | 3.78 | 10.33 <sup>b</sup> | 1.22 | 1.60 <sup>d</sup> | 0.25 |
| **Corolla Fusion Proportion** | 0.42 <sup>a</sup> | 0.07 | 0.64 <sup>b</sup> | 0.05 | 0.86 <sup>c</sup> | 0.08 | 0.65 <sup>b</sup> | 0.10 | 0.72 <sup>b</sup> | 0.08 | 0.21 <sup>d</sup> | 0.03 |
| **Lobe Length (mm)** | 5.48 <sup>a</sup> | 1.42 | 5.40 <sup>a</sup> | 0.70 | 2.77 <sup>b</sup> | 1.83 | 7.65 <sup>c</sup> | 0.66 | 4.08 <sup>a</sup> | 0.96 | 5.97 <sup>a</sup> | 0.11 |
| **Petal Length (mm)** | 9.53 <sup>a</sup> | 1.39 | 14.85 <sup>b</sup> | 1.04 | 18.41 <sup>c</sup> | 1.92 | 22.56 <sup>d</sup> | 2.98 | 14.40 <sup>b</sup> | 0.81 | 7.57 <sup>e</sup> | 0.22 |
| **Stamen Length (mm)** | 4.76 <sup>a</sup> | 0.38 | 10.80 <sup>b</sup> | 0.60 | 10.48 <sup>b</sup> | 1.00 | 10.94 <sup>b</sup> | 0.35 | 10.45 <sup>b</sup> | 0.57 | 6.15 <sup>c</sup> | 0.09 |
| **Anther Length (mm)** | 1.43 <sup>a</sup> | 0.10 | 1.59 <sup>b</sup> | 0.10 | 3.27 <sup>c</sup> | 0.19 | 1.86 <sup>d</sup> | 0.05 | 1.34 <sup>a</sup> | 0.15 | 5.75 <sup>e</sup> | 0.12 |
| **Ovary Diameter (mm)** | 1.78 <sup>a</sup> | 0.14 | 2.28 <sup>b</sup> | 0.19 | 5.30 <sup>c</sup> | 0.41 | 3.15 <sup>d</sup> | 0.18 | 1.37 <sup>e</sup> | 0.26 | 1.12 <sup>e</sup> | 0.06 |
| **Style Length (mm)** | 4.79 <sup>a</sup> | 0.53 | 7.90 <sup>b</sup> | 0.73 | 8.38 <sup>b</sup> | 0.66 | 12.74 <sup>c</sup> | 0.86 | 14.64 <sup>d</sup> | 1.47 | 5.45 <sup>a</sup> | 0.21 |
| **Nectar Volume (ul)** | 2.19 <sup>a</sup> | 0.48 | 6.86 <sup>b</sup> | 1.31 | 50.62 <sup>c</sup> | 3.76 | 32.89 <sup>d</sup> | 7.03 | 17.80 <sup>e</sup> | 3.84 | 0.00 <sup>a</sup> | 0.00 |

Sample sizes refer to the number of individuals measured (at least 3 flowers per individual). Significant differences among species are indicated by different letters ( $p < 0.001$ following Bonferri Correction, with ANOVA and Tukey's HSD). *Jaltomata* data from [42].

In addition to organ size and shape, our focal species also differ in the amount of nectar produced per flower (ANOVA *p* < 0.00001; Tukey HSD, *p* < 0.0001) (Table 2), as well as trichome type and density. In particular, the campanulate species *J. calliantha* and *J. dendroidea* as well as tubular *J. umbellata* produce significantly more nectar than either rotate *J. darcyana* or *J. sinuosa*. Finally, mature flowers differ in the types and density of trichomes on mature floral tissues. From simple to complex: *J. darcyana* has sparse simple (uniseriate) trichomes on both sepals and petals, *J. calliantha* dense simple trichomes on both sepals and petals, *J. umbellata* dense simple and dendritic (i.e. branched) trichomes on sepals but only dense simple ones on petals, *J. dendroidea* very dense dendritic trichomes on both sepals and petals, *J. sinuosa* very dense simple, dendritic and viscous glandular trichomes on sepals and dense simple and viscous glandular ones on petals, while *S. pimpinellifolium* has dense simple and non-viscous (i.e. non-secreting) glandular trichomes on both sepals and petals.

### The early sequence of floral development is similar across *Jaltomata* species

As expected based on their recent common origin (diverged <5 mya, [38]), the five focal species of *Jaltomata* share a common floral ground plan with each other as well as with the *Solanum* outgroup *S. pimpinellifolium* (from which they diverged ∼15-20 mya, [38]). All six species have very similar development throughout organ initiation, which corresponds to growth up to ∼1 mm bud diameter (stages A-F; Table 1). In all species, sepals are the first floral organs to emerge from the ∼0.15 mm diameter floral meristem, and do so in an asymmetric manner from the abaxial, to adaxial, to lateral sides (Figs. 2a, 3a, 5a-b, 6a, 7a). Once sepals within the outer whorl (within a flower) are qualitatively of similar size and trichome initiation is apparent, the former elongate from the base to form a partially fused structure (Figs. 2b, 3b, 4a, 5c, 6b, 7a-b). By ∼0.25-0.3 mm floral diameter, petals and stamens emerge almost simultaneously, but slightly asymmetrically (Figs. 2c, 3c, 4b-c, 5d-e, 6c-d, 7b), followed by gynoecial initiation by ∼0.5 mm diameter (Figs. 2f, 7c). Similar to sepals, petals initially emerge as free strap-like primordia, but begin to elongate from the congenitally fused region at the base (just above the stamen attachment point) by 0.7-0.8 mm diameter in all species (Figs. 2e, 3d, 4e, 5f-g, 6g, 7d). Simple (uniseriate) trichomes in *Jaltomata* species, and both simple and non-viscous glandular ones in *S. pimpinellifolium*, are first apparent on petals by ∼0.5 mm buds, followed by the emergence of dendritic petal trichomes in *J. dendroidea* and viscous glandular ones in *J. sinuosa* by 1 mm buds (Figs. 2e, 3d-e, 4d-e, 5g-h, 6f, 7e). Finally, the gynoecium is fused, and style elongation started, by ∼1 mm bud diameter in all species (Table 1; Figs. 2g, 3f, 4g-h, 5h-i, 6h-i). Because floral organs initiate at the same developmental age across species, observed differences in mature flowers are therefore not a result of differences in growth onset.

**Fig 2.**
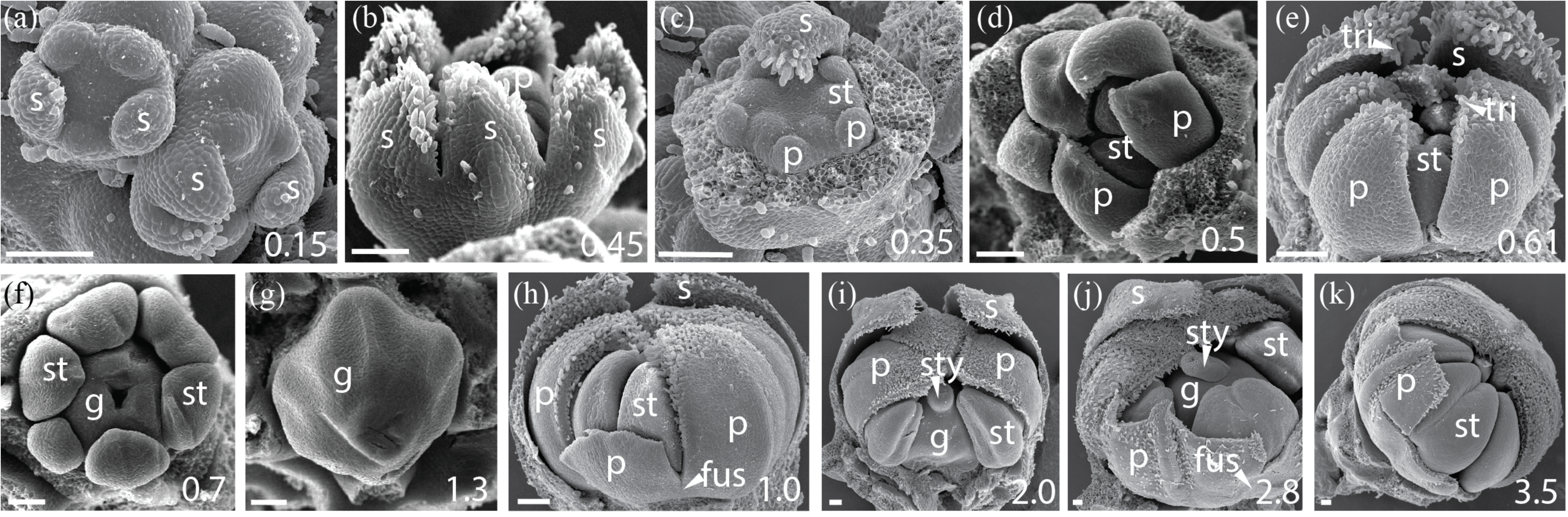
*Jaltomata darcyana* rotate flower development. (a,b) Stages A-C: Sepals develop asymmetrically from the abaxial side, and once of similar lengths, grow from a their base to form a fused tube. (c) Stage C: Removal of sepals reveals development of petal and stamen primordia. (d,e) Stage D: Petals grow freely but begin to develop trichomes. (f,g) Stages E-F: Carpels become fused during gynoecial development. (h-k) Stages F-G: Petals are congenitally fused at the base just above the stamen insertion point (h-i), but then become ‘superficially fused’ via interlocking trichomes along their lengths by ∼3 mm flower diameter (j-k). Scale bars = 100 um, and bud diameter is provided in bottom right panel. s = sepal, p = petal, st = stamen, tri = trichome, fus = corolla fusion, fil = anther filament, g = gynoecium, sty = style.

**Fig 3.**
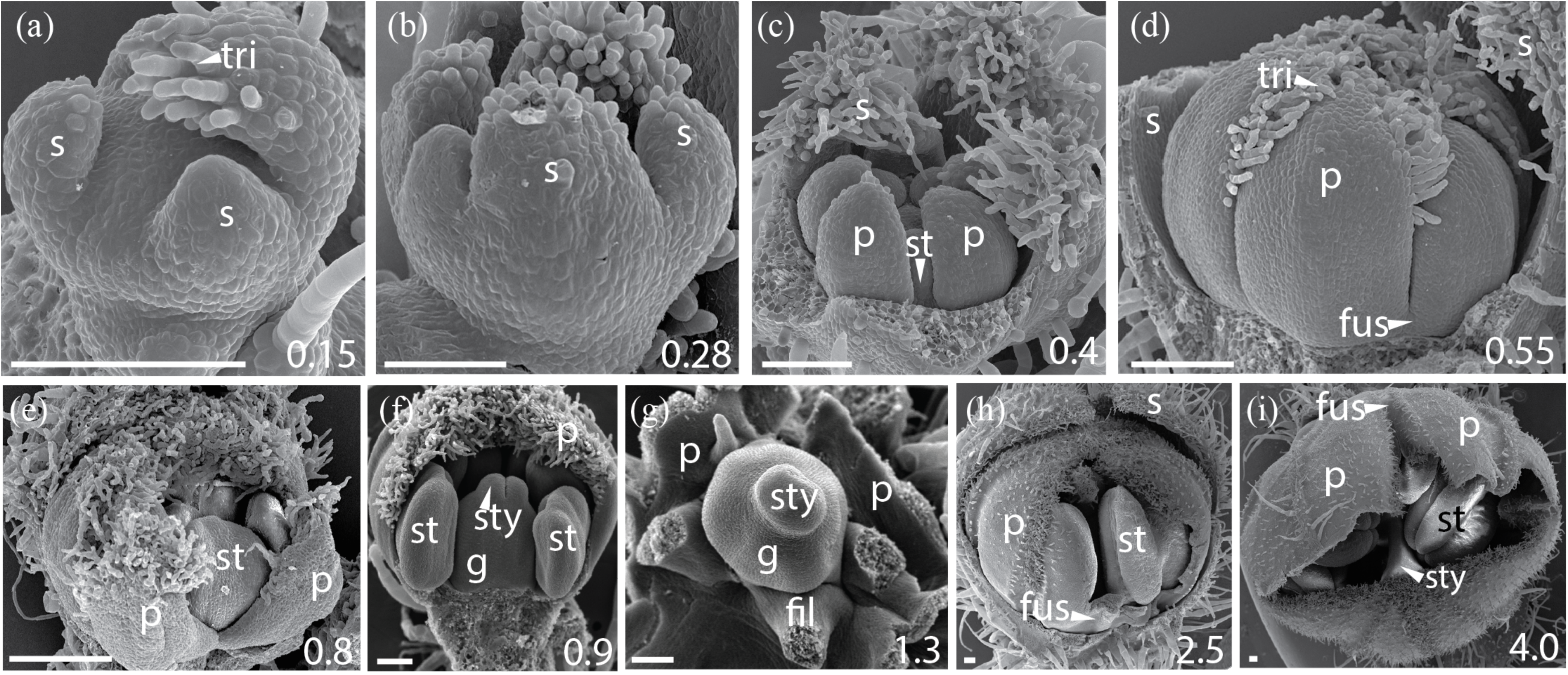
*Jaltomata sinuosa* rotate flower development. (a,b) Stages A-B: Sepals initiate in an asymmetric fashion from the abaxial side, and quickly fuse at the base. (c) Stage C: Five free petals develop around initiating stamen primordia. (d-e) Stage E: As stamens continue to mature, petals remain mostly free, but are fused at the base. (f-g) Stages F-G: Removal of mostly free petals reveals development of fused carpels. (h,i) Stage G: Opening of the corolla in 2.5-4 mm wide flowers shows a fused petal base from just above the stamen attachment point to about mid-way up the length of the corolla (i), and ‘superficial fusion’ of petals above the mid-point, via interlocking trichomes. Scale bars = 100 um, and bud diameter provided in bottom right corner. s = sepal, p = petal, st = stamen, tri = trichome, fus = corolla fusion, fil = anther filament, g = gynoecium, sty = style.

**Fig 4.**
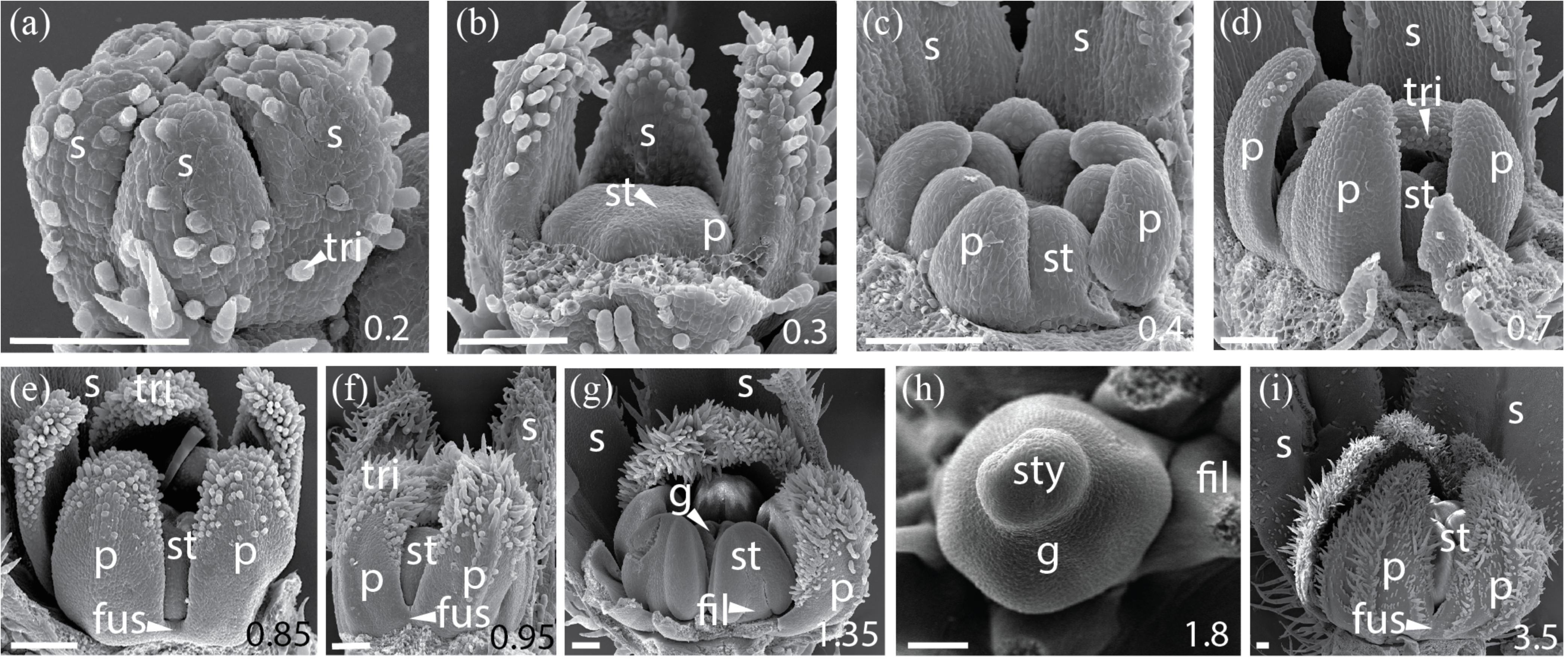
*Jaltomata calliantha* campanulate flower development. (a) Stage A: Free sepals extend through the growth of a basal primordium to form a lower fused tube. (b) Stage C: Opening of partially fused sepals reveals initiation of petal, followed by stamen, primordia. (c) Stage C: Free petals and stamens grow prior to initiation of the carpel primordia. (d-f) Stages D-E: Free petals develop trichomes on the adaxial side, becoming slightly fused just above the stamen attachment point during stamen thecae emergence. (g-h) Stages F-G: Fused carpel development. (i) Stage G: Petals remain only slightly fused above the stamen filament in 3.5 mm wide flowers. Scale bars = 100 um, and bud diameter is also provided in bottom right corner. s = sepal, p = petal, st = stamen, tri = trichome, fus = corolla fusion, fil = anther filament, g = gynoecium, sty = style.

**Fig 5.**
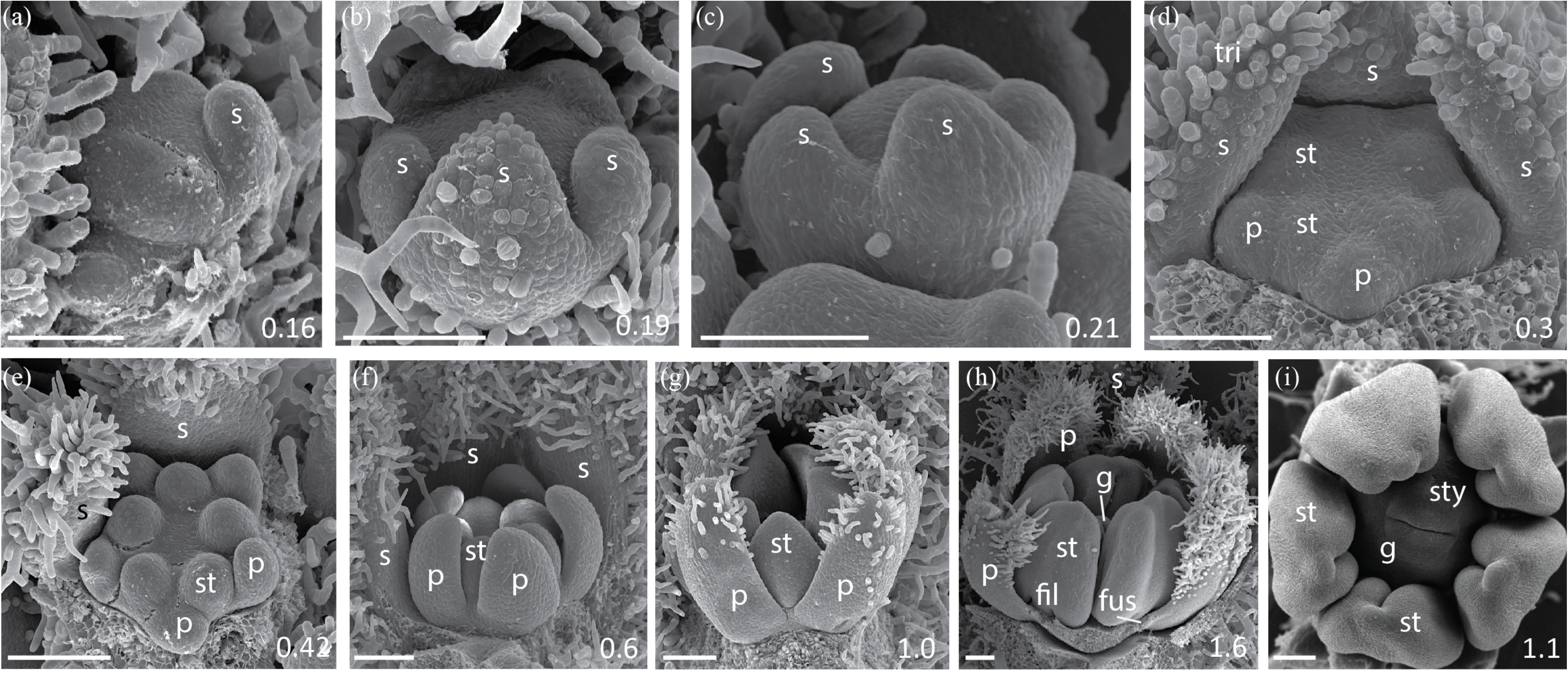
*Jaltomata dendroidea* campanulate flower development. (a,b) Stage A: Sepals develop asymmetrically from the abaxial side. (c) Stage A: Once equal in size, sepals extend through growth of the underlying meristem, giving rise to a fused tube. (d,e) Stage C: Partial removal of sepals reveals development of petals and stamens. (f-h) Stages D-G: Petals are largely free from early development to stamen thecae differentiation in 1.6 mm wide flowers, however, there is a small zone of fusion about the stamen attachment point. (i) Stage F: Carpels are partially fused above the developing ovary. Scale bars = 100 um, and bud diameter also provided in bottom right corner. s = sepal, p = petal, st = stamen, tri = trichome, fus = corolla fusion, fil = anther filament, g = gynoecium, sty = style.

**Fig 6.**
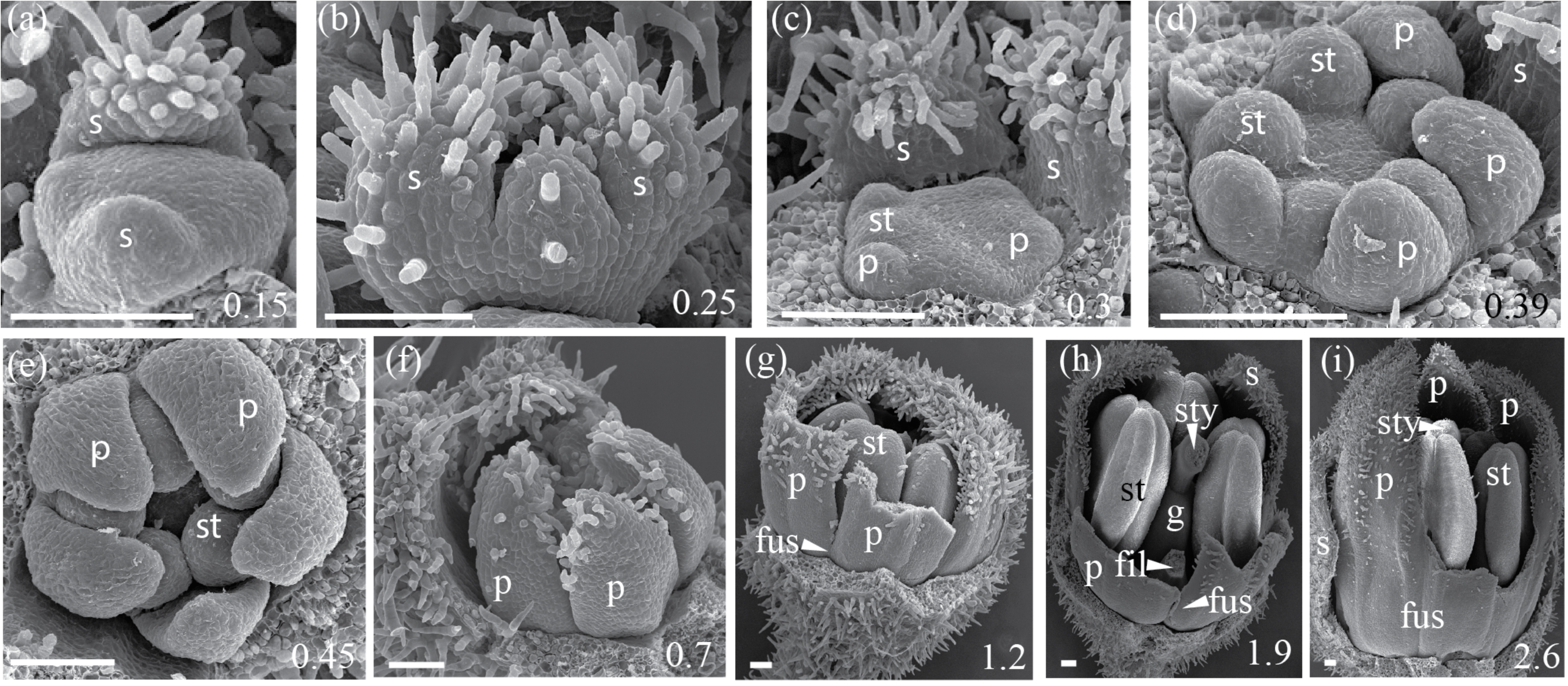
*Jaltomata umbellata* tubular flower development. (a) Stage A: Sepal development occurs asymmetrically, with the first sepal initiating on the abaxial side. (b) Stage B: Once all sepal primordia have expanded, organs become fused at the base to form a tube. (c,d) Stages C: Removal of partially fused sepals reveals the initiation of petal and stamen primordia in an asymmetric progression. (e-f) Stages C-D: Separate petal primordia elongate without fusion. (gi) Stages E-H: As stamens continue to mature, petals of ∼1-2.6 mm diameter flowers become increasingly fused above the stamen attachment point to form a corolla tube. By stage H (i), petals are longer than sepals. Scale bars = 100 um, and bud diameter also provided in bottom right corner. s = sepal, p = petal, st = stamen, tri = trichome, fus = corolla fusion, fil = anther filament, g = gynoecium, sty = style.

**Fig. 7.**
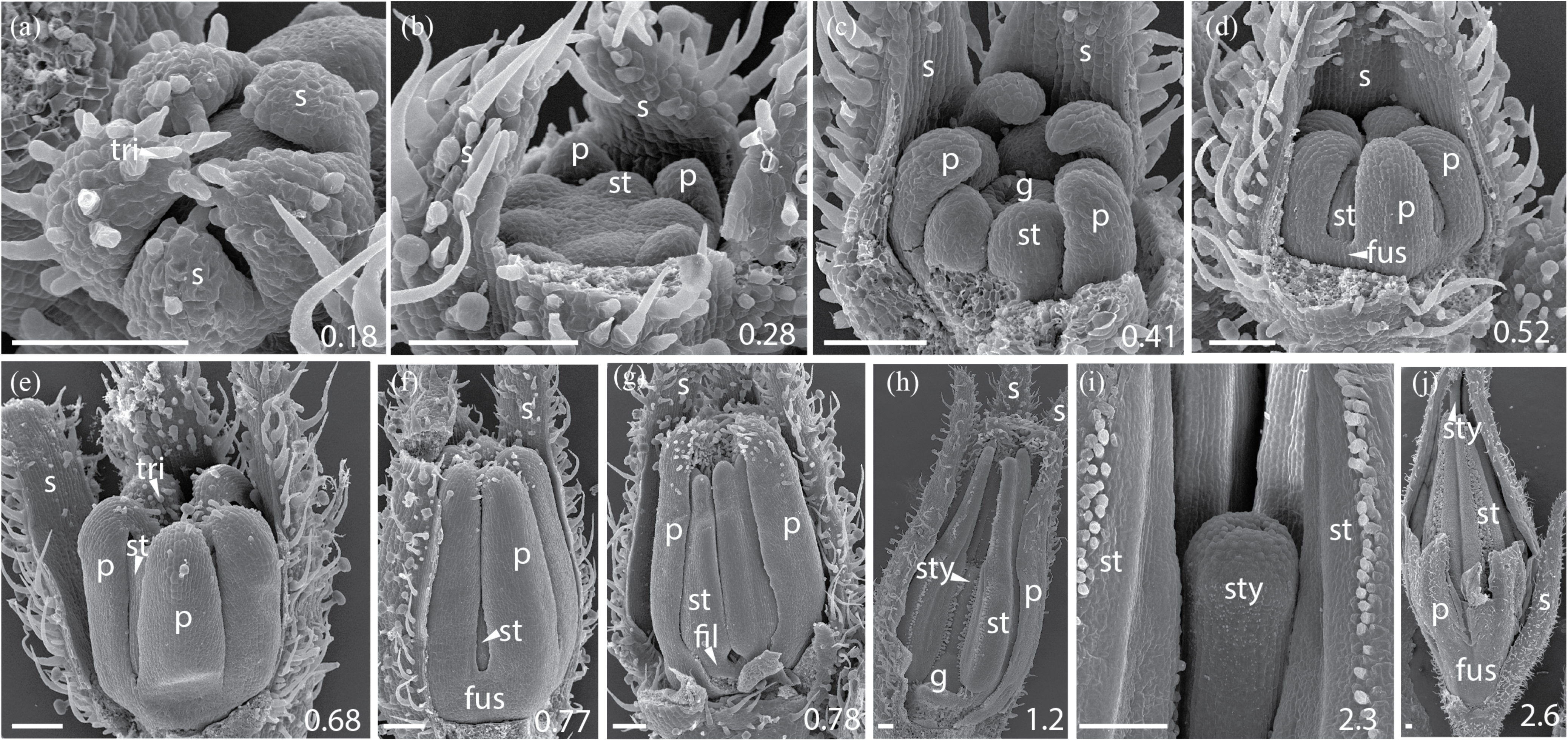
*Solanum pimpinellifolium* rotate flower development. (a) Sepals develop asymmetrically and fuse at their base to form a fused ring. (b,c) Removal of sepals reveals development of petals and stamens, followed by carpel development (c). (d-h) Petals are fused at the base above the stamen attachment point (visible in g) from early development (d), however, most of the petal margin is free. (i) Late carpel development. (j) At the 2 mm flower diameter stage, when anther thecae are well established, petals are mostly free, but remain partially fused at the base. Scale bars = 100 um, and bud diameter also provided in bottom right corner. s = sepal, p = petal, st = stamen, tri = trichome, fus = corolla fusion, fil = anther filament, g = gynoecium, sty = style.

**Fig. 8.**
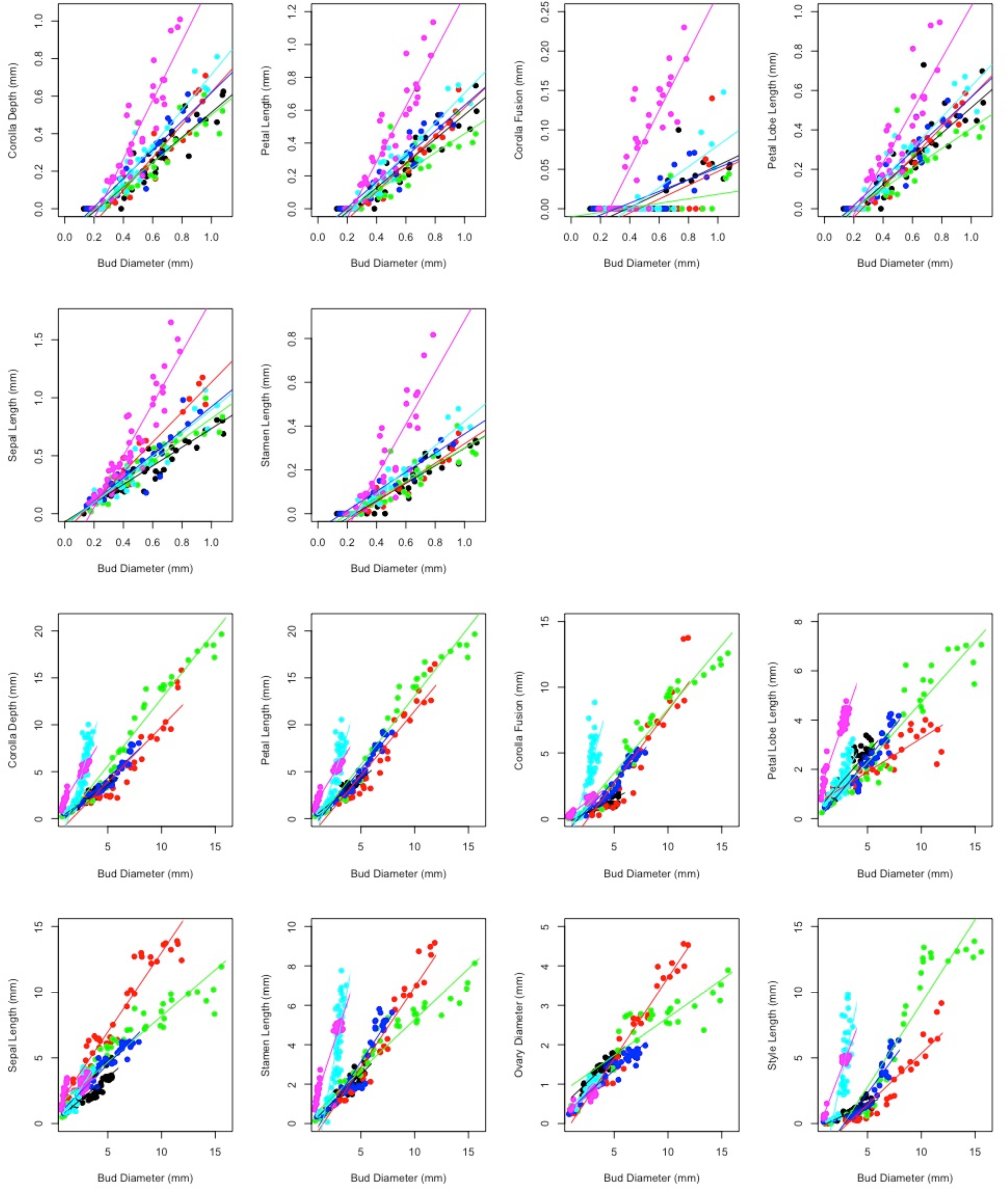
Growth trajectories for the focal floral organs and features. For all panels, the x-axis is bud diameter (mm) and the y-axis is the focal organ (mm). Top: growth rates during early stages (bud measurements during ‘stages A-F’); and Bottom: growth rates during mid- and late stages (bud measurements during ‘stages G-I’). Data points as well as regression lines of best fit are color coded by species. Slopes on log-transformed values are presented in Tables 4 and S3.

### Differential bud growth among species is apparent following organ initiation

Following the completion of floral organ initiation in ∼1 mm diameter buds, whole buds and their constituent organs continue to grow (including from the congenitally fused base for sepals and petals), anthers begin to mature with the appearance of two lobes or thecae, and trichomes mature on sepals and petals (“stage G” in Table 1). However, also during this stage, conspicuous differences in overall bud size as well as particular floral organs become apparent among species. Indeed, bud diameter begins to significantly differ among species during this mid-stage of floral development (ANOVA *p* < 0.0001), with *S. pimpinellifolium* buds remaining significantly smaller than those of all *Jaltomata* species except *J. umbellata* (Tukey HSD, *p* = 0.26 against *J. umbellata*, and *p* < 0.001 for all others), and *J. umbellata* remaining significantly smaller than all other *Jaltomata* species except *J. dendroidea* (Tukey HSD, *p* = 0.45 against *J. dendroidea*, and *p* < 0.05 for all others; Table 1). Despite this, corolla traits (corolla depth, corolla fusion, and petal length) do not significantly differ among species (ANOVA, all *p* > 0.1; Table S2), suggesting that their growth rates may be elevated in relation to whole bud growth within *S. pimpinellifolium* and *J. umbellata* (see below).

In addition to differences in bud and organ size among species, differences in petal trichome distribution also begin to emerge during stage G. In particular, trichomes along petal lobe edges (i.e. the portion of petals that are not part of the congenitally fused corolla tube) become interlocked, within *Jaltomata* species. Such interlocking trichomes extend from the region of congenitally fused corolla tube to petal tips, and appear to hold the edges of petal lobes together. We refer to this as ‘superficial -’ or ‘functional fusion’, as it does not result from fused petal tissue and petal lobes are easily separated once trichomes are pulled apart (Figs. 2h-k, 3f, h-i, 4g, i, 5h, 6g-h). This ‘superficial fusion’ is most apparent in buds of rotate *J. darcyana* and *J. sinuosa*, as well as campanulate *J. calliantha* and *J. dendroidea*, and only to a lesser extent within tubular *J. umbellata*. In particular, in rotate corollas of *J. darcyana* and *J. sinuosa*, the proportion of the congenitally fused corolla tube to region of ‘superficial fusion’ via interlocking trichomes remains similar throughout mid-stages; while in campanulate corollas of *J. calliantha* and *J. dendroidea* the congenitally fused region becomes proportionately larger during this time. In contrast to *Jaltomata*, petals in *S. pimpinellifolium* remain only slightly congenitally fused at the base, with no ‘superficial fusion’ via interlocking trichomes on lobe edges, during this stage (Fig. 7h).

During late stages of floral development, fully differentiated organs (especially petals) continue to grow and mature (stages H-I, Table 1) until petals open at anthesis (stage J, Table 1). Differences in bud diameter are especially pronounced during these late stages, with all species pairs significantly differing (ANOVA, *p* < 0.0001), except *J. umbellata* and S. *pimpinellifolium* (Tukey HSD, *p* = 0.99) during stage H, and *J. calliantha* and *J. dendroidea* (Tukey HSD, *p* = 0.47) and *J. umbellata* and S. *pimpinellifolium* (Tukey HSD, *p* = 0.99) during stage I (Table 1). At anthesis, trichomes providing ‘superficial fusion’ become unlocked, releasing petal lobes, while congenitally fused regions of the corolla remained fused in mature flowers.

Nectar secretion within *Jaltomata* flowers also begins during stages H-I of floral development, although there are some differences in timing among species. In *J. darcyana*, minute amounts of nectar are apparent in mid-day -1 day buds (i.e. the afternoon of the day before anthesis, during stage I); while for *J. calliantha*, *J. dendroidea*, and *J. sinuosa,* nectar is first apparent during the morning in 0 day buds (i.e. the morning of the day the flower opens, typically a few hours prior to full anthesis, during the very end of stage I); and in *J. umbellata*, nectar secretion is not apparent until the evening of day 0 flowers (after anthesis has occurred, during stage J), or until the morning of +1 day flowers (i.e. the day following anthesis, just prior to anther dehiscence in this species).

### Overall duration of floral development and absolute growth rates differ among species

We measured the duration of floral development in days to assess evidence that an extended growth period underpinned species floral differences (i.e. whether larger flowers grow for a longer period), as well as to calculate an average overall growth rate (bud diameter in mm per day) to assess potential accelerated growth of whole flowers.

The duration of floral development from 1 mm diameter buds onwards, which corresponds to completion of organ initiation in all species (i.e. starting in stage G, Table 1), was significantly different among species (Table 3). Floral development duration ranged from 14 days in outgroup *S. pimpinellifolium* to an average of 32.40 days in *J. dendroidea*. Among *Jaltomata* species, the ancestrally rotate flowers of *J. darcyana* (average 19.23 days) and derived tubular flowers of *J. umbellata* (average 18.62 days) had the shortest development times (ANOVA *p* < 0.00001; Tukey HSD against all other species, *p* < 0.00001; Tukey HSD against each other, *p* = 0.525). Similarly, overall average bud growth rate (bud diameter in mm per day) ranged from 0.185 mm/day in *J. umbellata* to 0.473 mm/day in campanulate *J. dendroidea* (Table 3).

**Table 3.** Average length of development, size of 0 day buds (just prior to anthesis), and overall bud growth rates for the five *Jaltomata* species and *S. pimpinellifolium*.

| Species | Length (days) |  | 0 Day Bud Diameter (mm) |  | Growth Rate (mm/day) |  |
| --- | --- | --- | --- | --- | --- | --- |
|  | Mean | SD | Mean | SD | Mean | SD |
| *J. darciana* (n=13) | 19.23<sup>a</sup> | 1.01 | 5.00<sup>a</sup> | 0.32 | 0.261<sup>a</sup> | 0.02 |
| *J. sinuosa* (n=11) | 28.55<sup>b</sup> | 0.93 | 7.16<sup>b</sup> | 0.45 | 0.251<sup>a</sup> | 0.02 |
| *J. calliantha* (n=10) | 28.80<sup>b</sup> | 1.03 | 12.13<sup>c</sup> | 0.36 | 0.422<sup>b</sup> | 0.02 |
| *J. dendroidea* (n=5) | 32.40<sup>c</sup> | 0.89 | 15.30<sup>d</sup> | 0.27 | 0.473<sup>c</sup> | 0.02 |
| *J. umbellata* (n=13) | 18.62<sup>a</sup> | 0.96 | 3.44<sup>e</sup> | 0.11 | 0.185<sup>d</sup> | 0.01 |
| *S. pimpinellifolium* (n=8) | 14.00<sup>d</sup> | 0.00 | 2.75<sup>f</sup> | 0.19 | 0.196<sup>d</sup> | 0.01 |

Bud diameter was measured every other day, from 1 mm buds to anthesis. Growth rate was determined by dividing bud diameter of day 0 buds by the length in days. Significant differences among species are indicated by different letters (following ANOVA and Tukey's HSD).

Because we also measured bud diameter while assessing duration of floral development, we compared these data with our measures from destructively sampled buds at different sizes to approximate how long (in days) each species remained at the different developmental stages (comprising mid- and late stages, Table 1). While this provides an imperfect estimate of the developmental time-course, it allowed us to assess whether there are evident qualitative differences between species. Based on this comparison, all *Jaltomata* species remained in stage F for approximately 2 days, but spent different amounts of time in mid- and each of the late stages (stages G-I) (Table 1). Rotate *J. darcyana* and tubular *J. umbellata* both spent fewer days in stage G than the other species. Interestingly, rotate *J. sinuosa* and campanulate *J. calliantha* followed similar time-courses, while campanulate *J. dendroidea* likewise had a similar time-course with the exception of a greatly extended duration of stage I.

### Relative floral organ growth rates differ among species

Given observed allometric differences in organ sizes among species (Table 2), we also calculated relative growth rates of individual floral organs. For instance, even if species do not differ in their overall absolute growth rate (diameter per day; Table 3), differences in the relative growth rates among floral organs (floral organ in mm per mm of bud diameter) could explain observed differences in mature flowers. Because we determined that overall bud growth rates do not differ among species until after completion of organ initiation (Table 1), we assessed relative organ growth rates during early stages separately from mid- and late stages (as outlined in Table 1).

We detected several differences among species in relative growth rates for particular floral organs, in both the early and the mid to late stage comparisons (Table 4, Fig. 8). First, during early stages, growth rates of all organs except the ovary and style are significantly elevated in rotate *S. pimpinellifolium* compared to rotate *J. darcyana* (all *p* < 0.00001, except *p* = 0.18 for ovary and *p* = 0.75 for style). Within *Jaltomata*, the growth rate for sepals is significantly elevated in campanulate *J. calliantha*, corolla fusion is significantly decreased in campanulate *J. dendroidea*, and as suggested by organ comparisons by stage (above), corolla depth is significantly elevated in tubular *J. umbellata* (Table 4). Comparing growth rates between campanulate *J. calliantha* and *J. dendroidea*, all corolla traits except fusion are significantly decreased in *J. dendroidea* (Table 4). Against *S. pimpinellifolium*, most organ growth rates are significantly decreased in *Jaltomata* species (indicative of general accelerated growth in *S. pimpinellifolium* starting in early stages), while only corolla growth rates are significantly decreased in *J. dendroidea* against both *J. darcyana* and *S. pimpinellifolium* (Table S3).

**Table 4.**
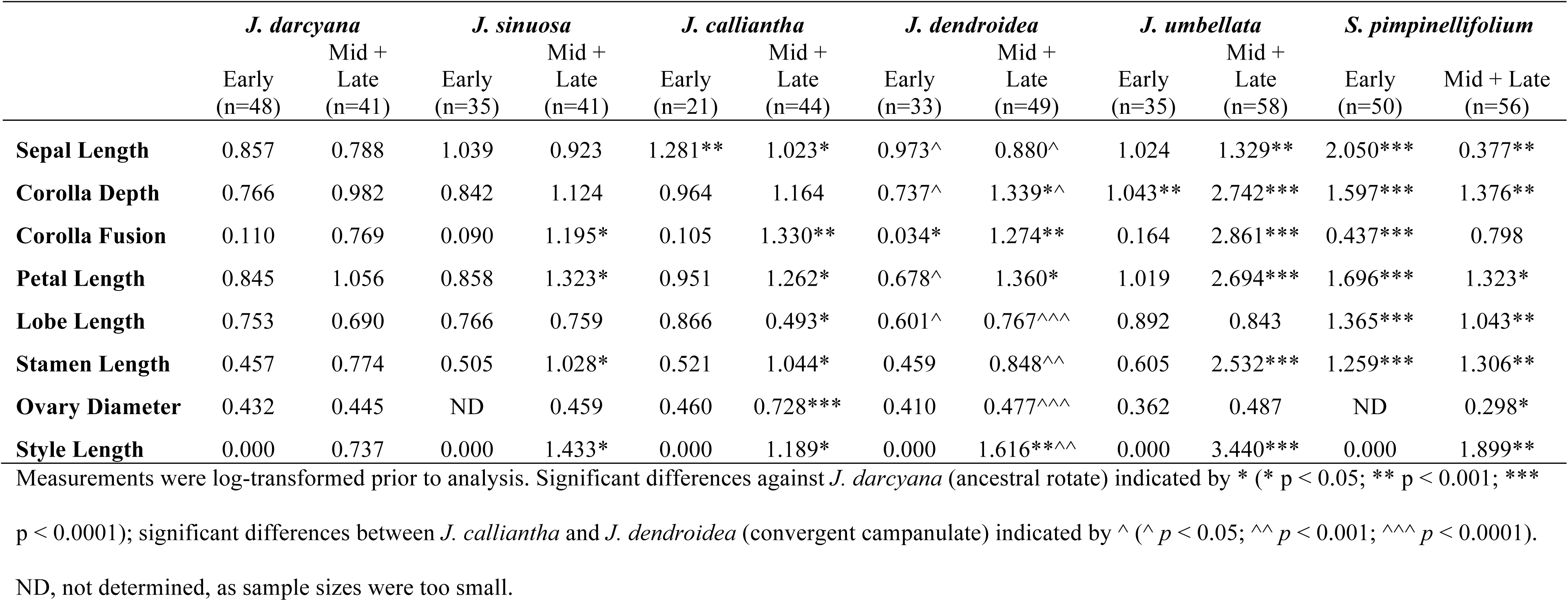
Relative growth rates for focal floral organs and features for the examined species, measured against bud diameter, for early and mid + late stages.

Within mid- and late stages of floral development, we detected even more differences in organ growth rates (Table 4, Fig. 8). Focusing on corolla growth within *Jaltomata*, corolla depth continues to be significantly elevated in *J. umbellata*; corolla fusion and petal length are significantly elevated in all other species; and lobe length is significantly decreased in *J. calliantha* (Table 4). In general, growth rates for corolla depth, corolla fusion, and petal length are highest in tubular *J. umbellata*. Growth rates for corolla fusion and petal length are similar between campanulate species (*J. calliantha* and *J. dendroidea*) and rotate *J. sinuosa*, however corolla depth and lobe growth rates are significantly elevated in *J. dendroidea* compared to *J. calliantha* (Table 4). Comparing against *S. pimpinellifolium*, corolla fusion is elevated and lobe length is decreased in rotate and campanulate *Jaltomata* species, and all corolla traits are elevated in tubular *J. umbellata*; while against both *J. darcyana* and *S. pimpinellifolium* all corolla traits are elevated in the other *Jaltomata* species, except lobe growth in *J. calliantha* (Table S3).

## Discussion

Heterochronic shifts during development are often considered a primary mechanism underlying phenotypic evolution [6]. Indeed, heterochrony appears to be a common mechanism underlying morphological diversification in numerous animal lineages [12], as well as in flowers, including shifts in both duration of floral development and growth rates leading to phenotypic shifts in descendent lineages [3-4, 17]. In this study, we examined floral ontogeny in five florally diverse species of *Jaltomata*, as well as a closely related outgroup that shares the ancestral rotate corolla form, to assess evidence for heterochrony contributing to observed variation in mature corolla forms and other floral traits. Overall, we found that the sequence of developmental events is consistent among examined species, but that differences in growth rate and duration of floral development are associated with different corolla morphologies. In particular, we found that early floral development (<1 mm bud diameter, corresponding to organ initiation) is very similar among all examined species, that differences between *Jaltomata* and *Solanum* are apparent soon after organ initiation (i.e. starting in stage G, Table 1), and that corolla trait differences associated with divergent corolla shapes among *Jaltomata* species first arise during mid-developmental stages. Elevated growth rates of corolla traits, combined with extended duration of floral development, lead to observed differences in mature floral traits among species. In particular, these developmental changes are heterochronic, with predominately accelerated growth rates and extended development duration (that is, peramorphic changes) associated with variation in mature corolla form.

### Extended duration of floral development and accelerated growth rates underlie derived tubular and campanulate mature corolla shapes in *Jaltomata*

One goal of this study was to assess whether derived tubular and campanulate corolla forms in *Jaltomata* arise due to heterochronic developmental shifts. In particular, we hypothesized that derived campanulate and tubular forms are elaborated versions of the ancestral rotate form, and therefore might result from peramorphic changes (i.e. longer duration of floral development or accelerated growth rates). Indeed, compared to ancestrally rotate flowers of *J. darcyana* (or indeed, more generally to *S. pimpinellifolium* or a composite of these two species), all derived *Jaltomata* species show peramorphic changes during floral development. First, all species except *J. umbellata* have significantly longer development duration (Table 3). Because floral organs initiated at the same developmental age across species, differences in the duration of floral development in derived forms result from changes in the timing of offset (i.e. growth period is extended). Second, growth rates for most of our measured floral traits are significantly elevated in *J. umbellata* compared to *J. darcyana*, including petal length and extent of corolla fusion (Table 4), even though *J. darcyana* and *J. umbellata* have similar development periods (19.23 days vs. 18.62 days, Table 3). Similarly, growth rates of most floral organs are significantly elevated in both campanulate species – including rates of petal length and extent of corolla fusion – in comparison to *J. darcyana* (Table 4), even though both also have longer development periods and faster overall growth rates in terms of bud diameter (Table 2). Thus, derived campanulate flowers in *J. calliantha* and *J. dendroidea* result from both accelerated organ growth rates and an extended development period, while derived tubular flowers in *J. umbellata* appear to result from accelerated growth rates of particular floral organs and organ components. These findings are consistent with other comparative ontogenetic studies examining floral morphology transitions associated with pollinator shifts, in which the derived floral form appears to be an elaborated version of the ancestral (e.g. [33]). For instance, peramorphic growth was found to underlie the elaborated nectariferous petal in the derived hummingbird pollinated *Delphinium* [33]. In contrast, in cases where the derived form appears to be a more juvenilized version, pedomorphic shifts predominantly explain these differences [4, 34-35].

In addition to changes associated with morphological differences between species, we also determined that nectar secretion dynamics appear to have undergone a combination of heterochronic shifts in more derived forms, including a delayed onset of the start of nectar secretion, as well as increased rates of nectar production. However, these shifts were found in rotate *J. sinuosa*, both campanulate *J. calliantha* and *J. dendroidea*, as well as tubular *J. umbellata*, suggesting that variation in nectar production is genetically and/or developmentally uncoupled from corolla morphology. Indeed, preliminary quantitative trait locus (QTL) mapping in a hybrid population generated from *J. sinuosa* and *J. umbellata* (Kostyun et al., unpublished) indicates that QTL for aspects of corolla morphology and nectar volume do not coincide. This is in contrast to studies in several other systems (reviewed in [48]) in which overlapping QTL were identified for floral shape and reward characters, although these cases as yet are unable to distinguish closely linked, but distinct, loci for corolla morphology and nectar traits, from actually pleiotropic loci.

### Alike mature corolla forms result from similar heterochronic changes during mid-stage ontogeny

Another goal of this study was to assess whether alike mature corolla shapes arise from shared or divergent developmental trajectories. Overall, we found that floral development was very similar during early and mid-stages within both species with rotate flowers, suggesting that these corolla morphologies may share a single evolutionary origin (i.e. are homologous). In contrast, species with campanulate flowers show several different types of changes, albeit in similar directions, supporting our hypothesis that the campanulate corolla morphologies in these two species are independently derived.

Floral ontogeny in general, and corolla development in particular, are very similar between rotate flowers of *J. sinuosa* and ancestrally rotate flowers of *J. darcyana*, until late stages when growth of petal length and corolla fusion become significantly elevated in *J. sinuosa* (Table 4). These species also share a similar overall bud growth rate, although *J. sinuosa* continues to grow for significantly longer (Table 3). These data indicate that the shared rotate floral morphology in these two species arises through a very similar developmental process, but peramorphic changes (both an extended development period and growth acceleration of particular floral organs) during late floral development lead to an overall increased size of *J. sinuosa* flowers, and a proportionately greater extent of corolla fusion (Table 2). These observations again suggest that the rotate flowers in these species share a single evolutionary origin.

In contrast, we determined that while both campanulate species have elevated corolla fusion and petal growth starting in mid-stage ontogeny (Table 4), they also significantly differ in perianth (sepal and petal) growth during early stages (Table 4), as well as in the duration of floral development (Table 3) and estimated duration of different stages (Table 1). Additionally, *J. calliantha* actually shows a significantly decreased growth rate (i.e. neoteny) for lobe length compared to *J. darcyana* during mid- and late stages (Table 4), whereas the growth rate of lobe length does not significantly differ between *J. dendroidea* and *J. darcyana*. These differences suggest that campanulate corollas in these two species are independently derived. Alternatively, these forms could share a single origin, but could have experienced subsequent developmental changes without large changes in the mature phenotype (i.e. developmental systems drift) during lineage divergence. In the absence of other data, we prefer the former hypothesis based on the phylogenetic distribution of these two species [37]; (Wu, Kostyun, Moyle, unpublished) (Fig. 1). Together, these observations agree with prior studies suggesting that different heterochronic changes contributing to similar mature floral phenotypes may actually be quite common; for instance, multiple distinct developmental shifts have been shown to produce flowers with the ‘selfing syndrome’ across lineages [3-4, 23] and among closely related populations and species [24].

### Heterochronic shifts as a first step to identifying specific mechanisms underlying floral divergence and convergence in *Jaltomata*

Our data suggest an important role for several different heterochronic developmental shifts in generating floral trait diversity among *Jaltomata* species, especially peramorphism (via growth acceleration and extended growth period), in shaping corolla diversity. If these heterochronic shifts are caused by relatively simple genetic changes, this could have contributed to the apparent rapid floral trait diversification within the genus (<5 my; [38]). Future work identifying the specific genetic and developmental mechanisms underlying these heterochronic shifts will provide a better understanding of the pace of phenotypic change, as well as likely candidate genes contributing to these developmental shifts. For instance, changes in the regulation of cell proliferation and/or cell expansion can lead to variation in overall floral size or size of particular floral organs, and several key candidate genes have been identified in *Arabidopsis* that function during these processes [23, 49]. Given our observed patterns of developmental heterochrony (i.e. extended floral development duration and accelerated growth), we anticipate that a combination of molecular heterochrony (e.g. growth promoting factors are expressed for longer) and heterometry (e.g. a higher amount of growth promoting gene products are produced) might underlie these shifts.

In addition to intrinsic (genetic) regulation of cell growth within floral organs, external signals or structures could also influence mature floral morphology [50-51]. For instance, the involvement of trichomes in regulating flower shape was recently reported in cotton (*Gossypium hirsutum*), in which the petal trichome gene *GhMYB-MIXTA-Like10 (GhMYBML10*) is strongly expressed at the point of petal overlap, resulting in trichome cross-linking that physically hold petals in place [46]. We identified similar ‘superficial corolla fusion’ via interlocking trichomes in both rotate and campanulate *Jaltomata* species examined here, although different growth dynamics during mid- and late-stages was observed between these forms. Although trichome cross-linking occurs in these species, it remains to be determined whether it is essential for normal corolla development (as in [46]) and if this phenotype is regulated by a *MYBML10* orthologue (including whether expression differs between rotate and campanulate vs. tubular *Jaltomata* forms).

The identification of specific developmental mechanisms and candidate genes will also clarify whether similar mature floral morphs result from shared evolutionary history or from convergent or parallel changes at the molecular level. In the *Jaltomata* species that we examined here, floral development (especially for corolla traits) in both rotate species was extremely similar until late stages, while in the campanulate species, we identified heterochronic shifts in the same direction but not in the same characters. Thus, it is likely that *J. darcyana* and *J. sinuosa* share their rotate form from common ancestry, while *J. calliantha* and *J. dendroidea* represent convergent or parallel evolution of the campanulate form. In this way, determining the developmental mechanisms underlying these heterochronic shifts can be used both to pinpoint how changes in existing floral development pathways lead to phenotypic variation, and to differentiate alternative evolutionary histories (common ancestry versus convergence) for similar mature floral forms. In particular, shared mutations would be more indicative of common descent rather than parallel evolution.

### Pollinator-mediated selection may have shaped *Jaltomata* floral diversification

Understanding how developmental trajectories differ between species with different mature floral traits also broadly informs the ways in which these traits are most able to respond to selection (e.g. whether developmental constraint restricts certain paths of trait evolution). Such inferences are especially relevant to floral trait evolution across the *Jaltomata* genus, which shows both high levels of floral trait divergence, as well as multiple putatively independent shifts to similar floral forms, within a relatively short timescale (<5mya, [38]). From an evolutionary perspective, such shifts are likely to have been shaped by pollinator behavior. Observations indicate that some species are predominantly visited by distinct pollinator functional groups (e.g. hymenopterans vs. hummingbirds, T. Mione, pers. comm.; J.L. Kostyun, unpublished), and these species have floral trait suites consistent with different pollinator syndromes [52]. In particular, hummingbirds have been observed visiting campanulate *J. calliantha* whose flowers have copious amounts of dilute nectar [53] as well as and tubular *Jaltomata viridiflora*, while hymenopterans have been observed visiting *J. sinuosa* and *Jaltomata repandidentata* (a close relative to *J. darcyana*; [37]) that have rotate corollas with relatively small amounts of concentrated nectar [54]. Because differential pollinator behavior often leads to reproductive isolation between lineages [55], understanding the underlying developmental basis of floral trait evolution can also reveal factors that could accelerate speciation in this rapidly-evolving and florally diverse system.

## Conclusions

As articulated by Darwin, vast diversity of form (including in flowers) can arise from changes in late developmental stages despite early developmental similarities that are rooted in common ancestry. One such type of developmental change is heterochrony, which is a shift in rate or timing in a descendant compared to its ancestor. By comparing floral ontogeny among five *Jaltomata* species and an outgroup, we determined that heterochronic shifts during mid- and late stages of floral development distinguish divergent corolla forms. In particular, two types of peramorphosis (differential growth acceleration and delayed offset leading to an extended development period) predominately explain these changes. Our data therefore support Darwin’s insight that even highly divergent mature floral traits result from modifications to initially similar structures. These relatively simple heterochronic shifts contribute to observed floral trait variation among *Jaltomata* species and, as such, could act as a mechanism allowing rapid floral trait diversification in this florally diverse system.

## Declarations

### Ethics approval and consent to participate

Not applicable.

### Consent for publication

Not applicable.

### Availability of data and material

Summaries of datasets used and/or analyzed during the current study are available within the Supplementary Materials, while raw data is available on Dryad [DOI to be added upon acceptance]. Seed material was obtained from Dr. Thomas Mione at Central Connecticut State University and the Tomato Genetics Resource Center at UC Davis – both of which followed all applicable regulations in obtaining the material.

### Authors’ contributions

All authors planned and designed the research, JLK performed the experiments, JLK and JCP analyzed the data, and all wrote the manuscript. All authors read and approved the final manuscript.

## Acknowledgements

We thank Armin Moczek and three anonymous reviewers for providing feedback on a previous draft of this manuscript, Barry Stein of the IU Electron Microscopy Center and Michele von

Turkovich of the UVM Microscopy Imaging Center for SEM and other instrument training, the IU Greenhouse staff for plant care, and Thomas Mione for sharing seed material.

## Additional files

**Additional file 1: Tables S1-S3. Table S1.** Accessions of the species used in the present study and their collection locations. All *Jaltomata* seed material provided by Dr. Thomas Mione at Central Connecticut State University (CCSU); *S. pimpinellifolium* seed provided by the Tomato Genetics Resource Center at UC Davis. **Table S2**. Summaries of average floral organ size at different stages of development across species. **Table S3**. Relative growth rates for floral organs for the examined species, measured against bud diameter, for early stages only, mid+late stages only, and overall development. Measurements were log-transformed prior to analysis. Significant differences against baseline indicated by * (* p < 0.05; ** p < 0.001; *** p < 0.0001), with black against J. darcyana, red against *S. pimpinellifolium* (alternate ancestral rotate), and blue against *J. darcyana* + *S. pimpinellifolium* (‘composite’ ancestral rotate); significant differences between *J. calliantha* and *J. dendroidea* (convergent campanulate) indicated by ^ (^ p < 0.05; ^^ p < 0.001; ^^^ p < 0.0001). Additional file 1 is in. xslx format.

**Additional file 2: Figure S1.** Measured morphological traits on mature flowers. Floral organs were measured on three flowers per individual, for at least three individuals per species, using hand-held digital calipers to the nearest 0.01 mm. Calyx diameter was measured as the widest planar distance across the calyx (outer most whorl); sepal length from the tip of the sepal to the center of the calyx; corolla diameter as the widest planar distance across the corolla (second whorl); corolla depth as the vertical distance from the base of the corolla to its highest point, petal length from the tip of the petal to the center of the corolla base; corolla fusion from the base of the corolla to the highest point of congenitally fused corolla; petal lobe length from the lowest point of non-fused petal edge to petal tip; stamen length from the base of the stamen (anther filament) to the tip of the anther; ovary diameter as the widest planar distance across the base of the ovary (inner most whorl); and style length from the base of the style to its tip (stigma). Additional file 2 is in .pdf format.

